# Ferroptosis is programmed by the coordinated regulation of glutathione and iron metabolism by BACH1

**DOI:** 10.1101/644898

**Authors:** Hironari Nishizawa, Mitsuyo Matsumoto, Tomohiko Shindo, Daisuke Saigusa, Hiroki Kato, Katsushi Suzuki, Masaki Sato, Yusho Ishii, Hiroaki Shimokawa, Kazuhiko Igarashi

## Abstract

Ferroptosis is an iron-dependent programmed cell death resulting from alterations of metabolic processes. However, its regulation and physiological significance remain to be elucidated. By analyzing transcriptional responses of murine embryonic fibroblasts exposed to the ferroptosis-inducer erastin, we found that a set of genes related to oxidative stress protection was induced upon ferroptosis. We further showed that the transcription factor BACH1 promoted ferroptosis by repressing the expression of a subset of erastin-inducible genes involved in the synthesis of glutathione or metabolism of intracellular labile iron, including *Gclm, Gclc*, *Slc7a11*, *Hmox1*, *Fth1, Ftl1*, and *Slc40a1*. Compared with wild-type mice, *Bach1*^-/-^ mice showed resistance to myocardial infarction, the seriousness of which was palliated by the iron-chelator deferasirox, which suppressed ferroptosis. Our findings suggest that ferroptosis is programmed at the transcriptional level to induce genes combating labile-iron-induced oxidative stress and executed upon disruption of the balance between the transcriptional induction of protective genes and accumulation of iron-mediated damage. BACH1 is suggested to control the threshold of ferroptosis and to be a therapeutic target for palliating myocardial infarction.

## Introduction

Ferroptosis is a new form of programmed cell death caused by the iron-dependent accumulation of lipid hydroperoxide (Dixon et al., 2012, Yang et al., 2014). As a pathological cell death, ferroptosis causes various oxidative stress-related diseases, including ischemia-reperfusion injury (Baba et al., 2018, Fang et al., 2019, Gao et al., 2015, Linkermann et al., 2013, Linkermann et al., 2014) and neurodegenerative diseases (Chiang et al., 2017, Di Domenico et al., 2017). Ferroptosis also contributes to tumor suppression as a response induced by p53 and is important for organisms in preventing cancer (Jiang et al., 2015, Kim et al., 2016, Viswanathan et al., 2017, Yang et al., 2014). Considering the involvement of lipid hydroperoxide, ferroptosis may be executed at the edge of the oxidative stress response. Therefore, ferroptosis may be a regulated process involving the oxidative stress response. However, the regulatory mechanism underlying ferroptosis has yet to be elucidated in full.

BTB and CNC homology 1 (BACH1) is a heme-binding transcription factor required for the proper regulation of the oxidative stress response and metabolic pathways related to heme and iron (Ogawa et al., 2001, Sun et al., 2004, Suzuki et al., 2004). BACH1 represses *Hmox1* encoding heme oxygenase-1 (HO-1), *Fth1* and *Ftl1* encoding ferritin proteins, *Gclm* and *Gclc* encoding glutamate-cysteine ligase modifier and catalytic subunits, and other genes involved in the oxidative stress response (Hintze et al., 2007, Sun et al., 2002, Warnatz et al., 2011). We hypothesized that BACH1 might regulate ferroptosis by inhibiting the expression of these genes. In addition, since BACH1 is involved in the exacerbation of various diseases involving oxidative stress, such as ischemic heart disease (Yano et al., 2006), hyperoxic lung injury (Ito et al., 2017), trinitrobenzene sulfonic acid-induced colitis (Harusato et al., 2013), nonalcoholic steatohepatitis (Inoue et al., 2011), and spinal cord injury (Kanno et al., 2009), BACH1 may exacerbate the severity of these diseases through ferroptosis.

To understand the regulatory mechanism underlyning ferroptosis, we analyzed the transcriptome response in ferroptotic cells with RNA sequencing (RNA-seq). We also examined whether or not BACH1 was involved in the regulation of ferroptosis by comparing ferroptosis and the expression of ferroptosis-induced genes between wild-type (WT) and *Bach1*^-/-^ murine embryonic fibroblasts (MEFs). Furthermore, we assessed the influence of BACH1 and ferroptosis on the severity of acute myocardial infarction (AMI) in model mice. We found that BACH1 promoted ferroptosis by directly repressing genes involved in the synthesis of glutathione (GSH) and sequestration of free labile iron. BACH1 also increased the severity of AMI, which was mitigated by the iron chelator deferasirox (DFX). Our findings highlight the coordinated transcriptional response and its regulation by BACH1 upon ferroptosis.

## Results

### Transcriptomic alterations in ferroptotic cells

Changes in metabolic and biological processes occur in ferroptotic cells (Shimada et al., 2016, Yang et al., 2016). To determine the changes in transcriptome upon ferroptosis, we treated MEFs with erastin, a class I ferroptosis inducer (Dixon et al., 2012, Yang et al., 2014), and carried out RNA-seq analyses. Genes related to oxidative stress and iron metabolism showed significant inductions in their expression (Fig 1A). Some of these genes are known to possess inhibitory effects on ferroptosis (Stockwell et al., 2017). Therefore, ferroptosis accompanies the induction of genes that can restrict the execution of ferroptosis.

**Figure 1.**
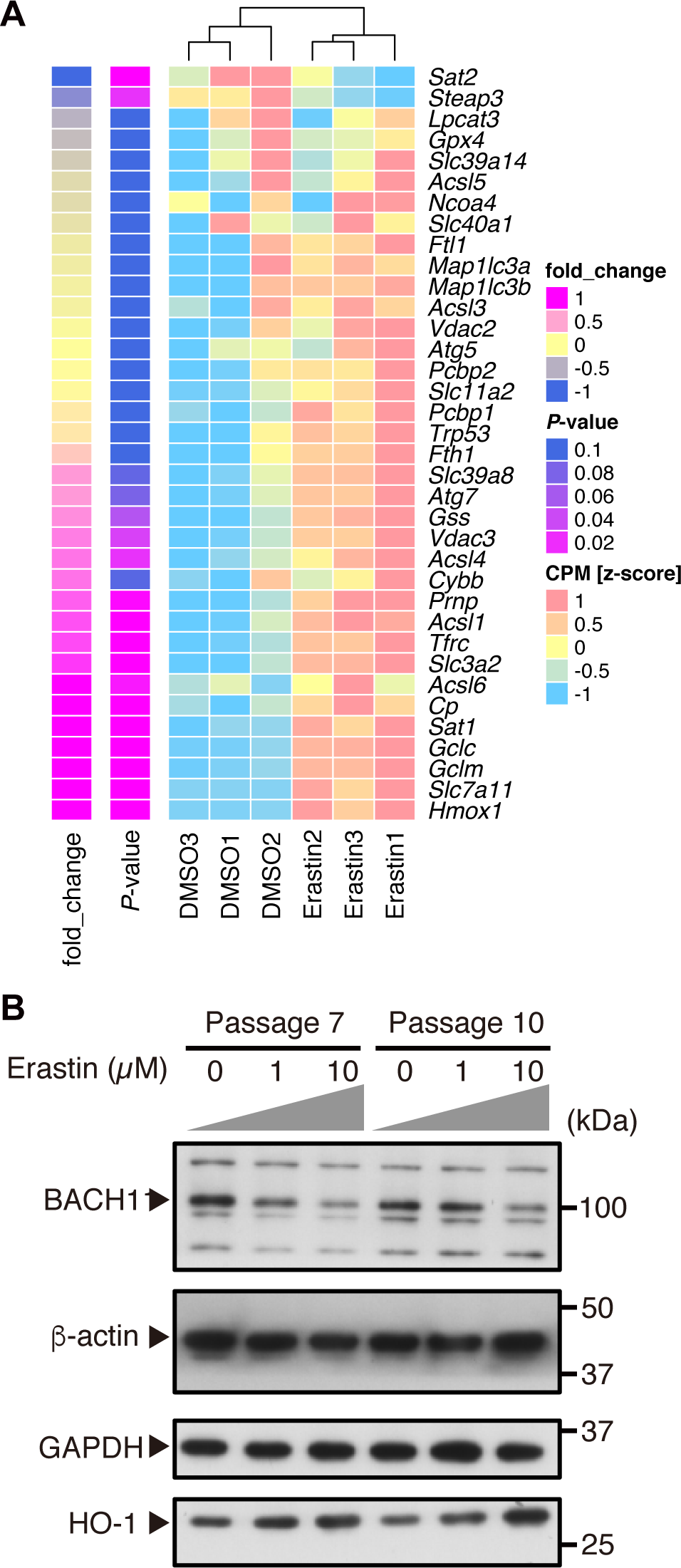
Regulartory genes of ferroptosis are upregulated with decreasing BACH1 protein at the induction of ferroptosis. A RNA-seq was performed in WT MEFs (9th passage: P9) with only DMSO (DMSO group) or DMSO + 3 µM Erastin (Erastin group) for 24 hrs. A heat map of gene expression profiles shows the genes registered to map04216 (Ferroptosis pathway) of Kyoto Encyclopedia of Genes and Genomes (KEGG) pathway map. The genes were arranged from the bottom in the order of fold change of Erastin group to DMSO group. n = 3 per group. B Western blotting for BACH1, HO-1, β-actin, and GAPDH of WT MEFs (P7, P11) exposed to erastin for 12 hrs. Data information: In (A), *P*-value by the differential expression analysis performed on edge R.

Among the induced genes, *Hmox1* encoding HO-1 is reported to be associated with ferroptosis (Kwon et al., 2015, Sun et al., 2016) and is a well-known target of BACH1 (Kitamuro et al., 2003, Sun et al., 2002). *Slc7a11* encodes a component of system x_c_^-^(cystine/glutamine transporter) (Sato et al., 2005, Sato et al., 2000) and a well-known regulator of ferroptosis (Jiang et al., 2015). *Gclm* and *Gclc* encode glutamate-cysteine ligase modifier and catalytic subunits (Fan et al., 2012, Telorack et al., 2016), both considered to suppress ferroptosis by GSH synthesis (Miess et al., 2018, Stockwell et al., 2017). These genes for the pathway of GSH synthesis are also considered to be targets of BACH1 (Warnatz et al., 2011). Indeed, the amount of BACH1 protein was decreased in MEFs exposed to erastin, which was accompanied by the induction of *Hmox1* (Figs 1B and EV1). With the reduction in BACH1 protein, the production of its mRNA was induced (Fig EV1), suggesting the presence of feedback regulation of BACH1.

These observations suggest that, when cells are exposed to erastin, the expression of genes that counteract ferroptosis is induced in part by a reduction in BACH1 protein and that the amount or activity of BACH1 and the kinetics of its feedback regulation may influence ferroptosis by suppressing this counteracting subprogram of ferroptosis.

### BACH1 promotes ferroptosis

To clarify whether or not BACH1 regulates ferroptosis, we treated WT and *Bach1*^-/-^ MEFs with erastin, stained them with propidium iodide (PI) and annexin V, and compared the cell death by a flow cytometry analysis (Fig EV2). *Bach1*^-/-^ MEFs showed less cell death in response to erastin than WT cells (Fig 2A and B). When the erastin-treated MEFs were observed with a transmission electron microscope, shrunken mitochondria, which are characteristic of ferrotosis (Dixon et al., 2012), were confirmed in both WT and *Bach1*^-/-^ MEFs (Fig 2C). The cell death in our experiments was inhibited by the iron chelator deferoxamine (DFO) (Fig 2D-F), confirming that this death was ferroptosis. These results showed that BACH1 promoted ferroptosis in MEFs.

**Figure 2.**
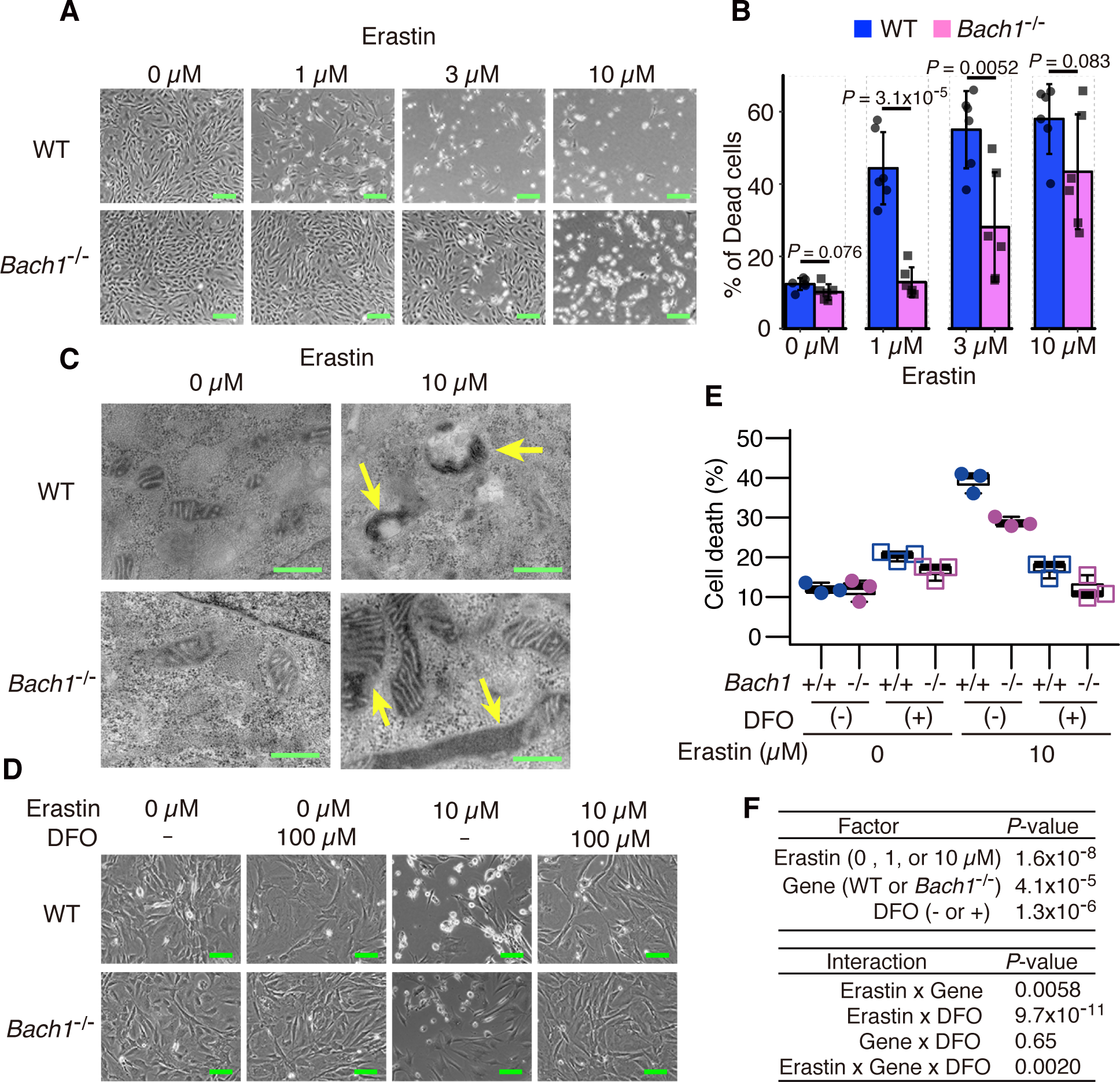
BACH1 promotes ferroptosis. A, B Optical microscope image (A) and Quantification of cell death by flow cytometer (B) of WT and *Bach1^-/-^* MEFs (11th passage: P11) exposed to erastin for 24 hrs. Scale bars in (A) represent 100 µm. C Transmission electron microscope image of WT and *Bach1^-/-^* MEFs (P9) exposed to erastin for 10 hrs. Arrow: shrunken mitochondria. Scale bars represent 500 nm. D-F Optical microscope image (D) and Quantification of cell death by flow cytometer (E,F) of WT and *Bach1^-/-^* MEFs (P12) exposed to erastin and DFO for 24 hrs. (F) is statistical analysis results of (E). Scale bars in (D) represents 200 µm Data information: (A, B, D, and E) are representative of three independent experiments. Error bars of (B) represent standard deviation. The box and whisker plots of (E) show the 25th and 75th percentile quartiles and median values (center black line) and maximum and minimum values of the data. *P*-value of (B) by *t*-test. *P*-value of (F) by three-way ANOVA.

It should be noted that the difference in cell death between WT and *Bach1*^-/-^ MEFs became smaller as the dose of erastin increased (Fig 2A and B). This may be because even *Bach1*^-/-^ MEFs lost their resistance to ferroptosis under high doses of erastin. This suggests that the function of BACH1 is more meaningful for restricting ferroptosis under low-stress conditions. Therefore, the reduction in BACH1 protein may be part of the early ferroptosis program, and BACH1 may set the threshold for ferroptosis. Execution of ferroptosis may be determined by the basal amount of BACH1 and how rapidly it is degraded in response to ferroptosis inducers.

### BACH1 represses the expression of genes involved in the GSH synthesis pathway

BACH1 may decrease GSH by repressing the expression of genes involved in the pathway of GSH synthesis. To investigate this possibility, we measured the intracellular GSH concentrations in WT and *Bach1*^-/-^ MEFs. The amount of GSH was significantly higher in *Bach1*^-/-^ MEFs than in WT cells (Fig 3A), suggesting that BACH1 promoted ferroptosis by reducing GSH within cells.

**Figure 3.**
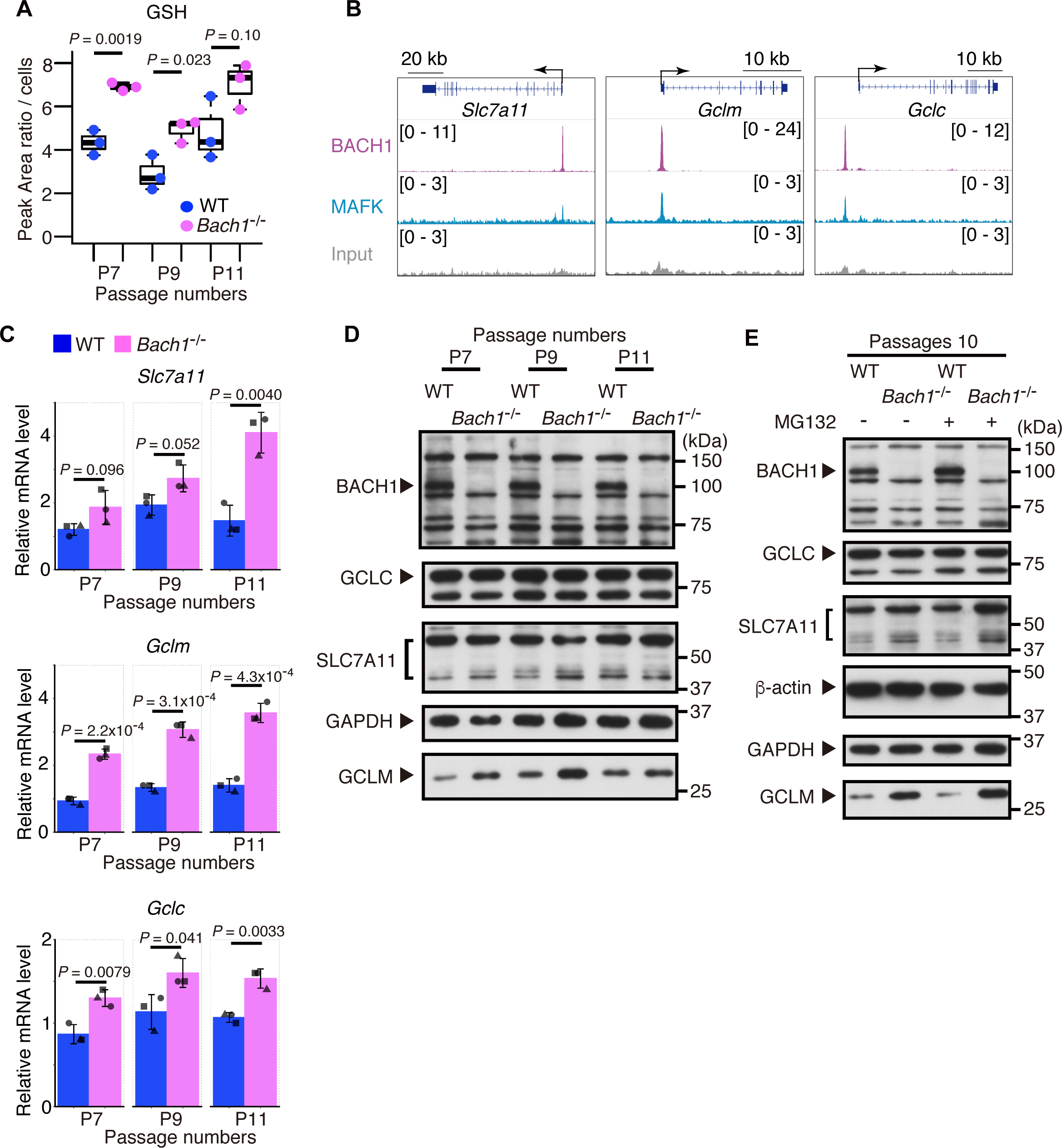
BACH1 decreases GSH by repressing *Slc7a11*, *Gclm*, and *Gclc* expression. A Intracellular concentration of GSH in WT and *Bach1^-/-^* MEFs (7th, 9th, and 11th passage: P7, P9, and P11) by UHPLC/MS/MS. B ChIP-seq analysis of the binding of BACH1, MAFK for gene regions of *Slc7a11*, *Gclm*, and *Gclc* in M1 cells. C qRT-PCR analysis for *Slc7a11*, *Gclm*, and *Gclc* mRNA relative to *Actb* mRNA in WT and *Bach1^-/-^*MEFs (P7, P9, P11). n = 3 of independent rots of MEFs per genotype. D Western blotting for BACH1, SLC7A11, GCLM, GCLC, and GAPDH of WT and *Bach1^-/-^* MEFs (P7, P9, P11). E Western blotting for BACH1, SLC7A11, GCLM, GCLC, b-actin and GAPDH in WT and *Bach1^-/-^* MEFs (P10) exposed to 25 µM MG132. Data information: The box and whisker plots of (A) show the 25th and 75th percentile quartiles and median values (center black line) and maximum and minimum values of the data. Error bars of (C) represent standard deviation. *P*-value of (A,C) by *t*-test.

**Figure 4.**
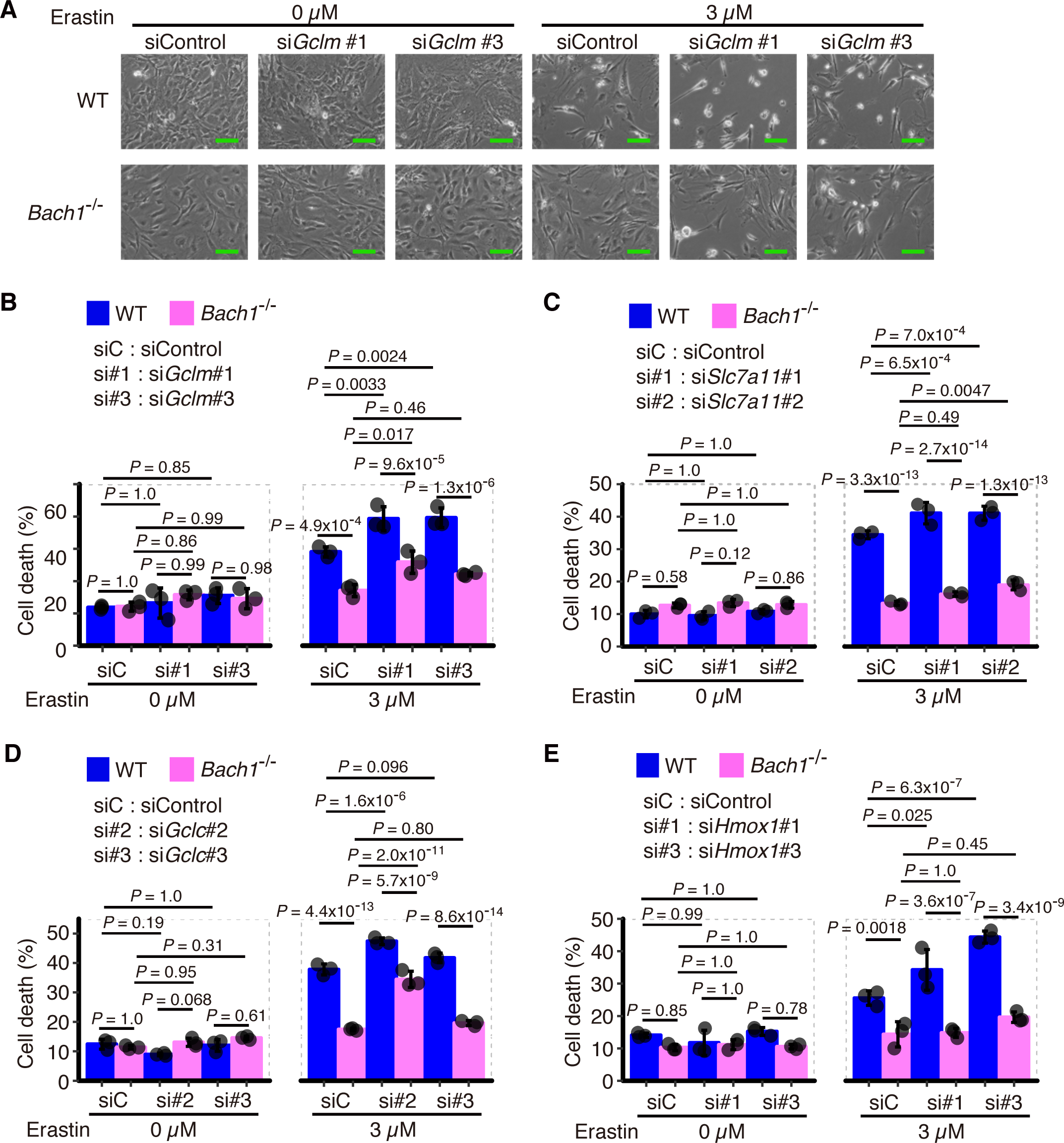
*Gclm*, *Slc7a11*, *Gclc*, and *Hmox1* repress ferroptosis. A-E siRNA was transfected to WT and *Bach1*^-/-^ MEFs (5th or 6th passage). After 24 hrs, MEFs were exposed to erastin for 24 hrs. Optical microscope image (*A*) and Quantification of cell death by flow cytometer (*B-E*). Scale bars in (A) represent 100 µm. Data information: Error bars of (B-E) represent standard deviation. *P*-value by Tukey’s test after three-way ANOVA.

By revisiting our previous data of chromatin immunoprecipitation with sequencing (ChIP-Seq) of BACH1 in mouse myeloblast M1 cells (Ebina-Shibuya et al., 2017, Ebina-Shibuya et al., 2016), we found peaks of BACH1 and its partner MAFK in the regulatory regions of genes encoding molecules for glutathione synthesis, including *Gclm*, *Gclc*, and *Slc7a11* (Fig 3B). Furthermore, by comparing the expression of these genes in WT and *Bach1*^-/-^ MEFs by quantitative polymerase chain reaction (qPCR), the expression of all of these genes was confirmed to be higher in *Bach1*^-/-^ MEFs than in WT cells (Fig 3C). These results suggested that BACH1 bound to the regulatory regions of these genes to repress their expression.

A comparison of the protein amounts of SLC7A11, GCLM, and GCLC in MEFs by Western blotting revealed that more GCLM protein was present in *Bach1*^-/-^ MEFs than in WT cells (Fig 3D and E). Although the amounts of SLC7A11 protein were similar in WT and *Bach1*^-/-^ MEFs (Fig 3D), more SLC7A11 protein was present in *Bach1*^-/-^ MEFs than in WT cells when they were treated with proteasome inhibitor MG132 (Fig 3E). These observations suggest that the amount of SLC7A11 protein is further tuned by proteasomal-mediated degradation. There were no marked differences in the amount of GCLC protein with or without MG132 (Fig 3D and E). BACH1 may affect the expression of GCLC protein under certain circumstances. Given these results, we surmised that BACH1 decreased the amount of GSH in part by repressing the expression of *Gclm* and *Slc7a11*.

### BACH1 promotes ferroptosis by altering GSH

We next examined whether or not the resistance of *Bach1*^-/-^ MEFs against ferroptosis was actually dependent on the increased expression of the genes involved in the GSH synthesis pathway. Although it is not always statistically significant, knockdown of any of *Slc7a11, Gclm,* and *Gclc* resulted in slight but reproducible increases in ferroptosis in both WT and *Bach1*^-/-^ MEFs (Figs 4A-D and EV3A, B. Appendix Fig S1A-C). These results show that the genes involved in the GSH synthesis pathway have inhibitory effects against ferroptosis and suggest that BACH1 promotes ferroptosis by repressing their expression.

We next examined the effect of knockdown of *Hmox1*. WT MEFs became more sensitive to ferroptosis by knockdown of *Hmox1* than cells with control knockdown (Figs 4E and EV3C. Appendix Fig S1D). We thus concluded that HO-1 works as an inhibitor of ferroptosis under our experimental conditions. However, the effect of HO-1 to accelerate ferroptosis has also been reported (Fang et al., 2019, Kwon et al., 2015). The function of HO-1 in ferroptosis might differ depending on the situations of cells.

Importantly, knockdown of *Slc7a11, Gclm, Gclc,* or *Hmox1* did not decrease the observed differences in ferroptosis between WT and *Bach1*^-/-^ MEFs (Figs 4A-E and EV3A-C. Appendix Fig S1A-D). These results suggest that the role of BACH1 in promoting ferroptosis depends on the repression of multiple genes involved in ferroptosis.

### BACH1 accelerates ferroptosis by suppressing labile iron metabolism

To explore other target genes of BACH1 in the regulation of ferroptosis, we examined genes involved in the regulation of iron metabolism (*Fth1*, *Ftl1*, *Slc40a1*, *Tfrc*, *Mfn2*, and *Fxn*), heavy metal stress (*Mt1*), and lipoperoxidation (*Gpx4*). Some of these genes were upregulated in response to erastin (see Fig 1A). Among these genes, ferritin genes (*Fth1* and *Ftl1*) and the ferroportin gene (*Slc40a1*) were dramatically upregulated in *Bach1*^-/-^ MEFs (Fig 5A), and binding peaks of BACH1 and MAFK were observed near their regulatory regions (Fig 5B). In contrast, the expression of *Tfrc*, *Mfn2*, *Fxn*, *Mt1*, and *Gpx4* was only mildly increased in *Bach1*^-/-^ MEFs (Fig EV4A). There were no strong binding peaks of BACH1 or MAFK in the regulatory regions of these genes (Fig EV4B). Considering that both ferritin and ferroportin reduce the availability of free labile iron and are known to inhibit ferroptosis (Geng et al., 2018, Wang et al., 2016), these results suggest that BACH1 promotes ferroptosis by repressing the transcription of ferritin and ferroportin genes. These findings, along with the regulation of GSH synthesis pathway by BACH1, suggest that BACH1 accelerates ferroptosis by decreasing the intracellular activity of GSH and increasing the oxidative activity of labile iron (Fig 5C).

**Figure 5.**
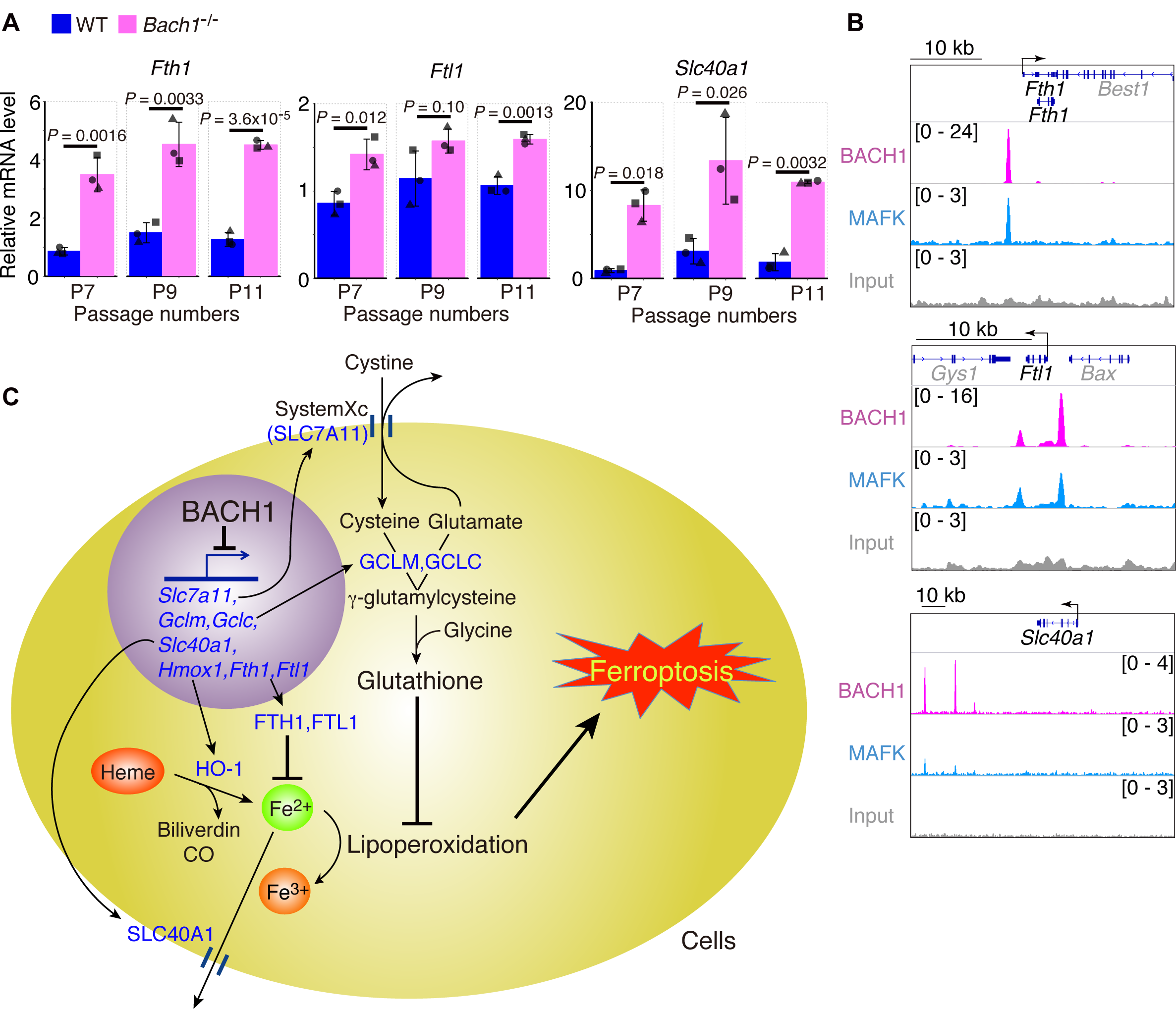
BACH1 represses transcription of genes of ferritin and ferroportin. A qRT-PCR analysis for *Fth1*, *Ftl1*, and *Slc40a1* mRNA relative to *Actb* mRNA in WT and *Bach1^-/-^* MEFs (7th, 9th, and 11th passage: P7, P9, and P11). n = 3 of independent rots of MEFs per genotype. B ChIP-seq analysis of the binding of BACH1, MAFK for gene regions of *Fth1*, *Ftl1*, and *Slc40a1* in M1 cells. C Conseptual diagram. Data information: Error bars of (A) represent standard deviation. *P*-value of (A) by *t*-test.

### BACH1 aggravates acute myocardial infarction by promoting ferroptosis

Finally, we tried to examine whether or not the promotion of ferroptosis by BACH1 is involved in pathological changes *in vivo*. As there are several reports showing that ferroptosis is involved in ischemia-reperfusion injury in the heart (Baba et al., 2018, Fang et al., 2019, Gao et al., 2015), we used an AMI model based on left anterior descending coronary artery (LAD) ligation (Abarbanell et al., 2010, Shindo et al., 2016) (Fig 6A). In this model, *Bach1*^-/-^ mice showed less severe injuries than WT mice as judged by the post-operative survival rate and an evaluation of the cardiac function with echocardiography (Figs 6B, C and EV5A-C. Movie EV1A-D). The infarct area on pathological specimens was also smaller in *Bach1*^-/-^ mice than in WT mice (Fig 6D and E). These results suggest that BACH1 exacerbates the pathology of AMI.

**Figure 6.**
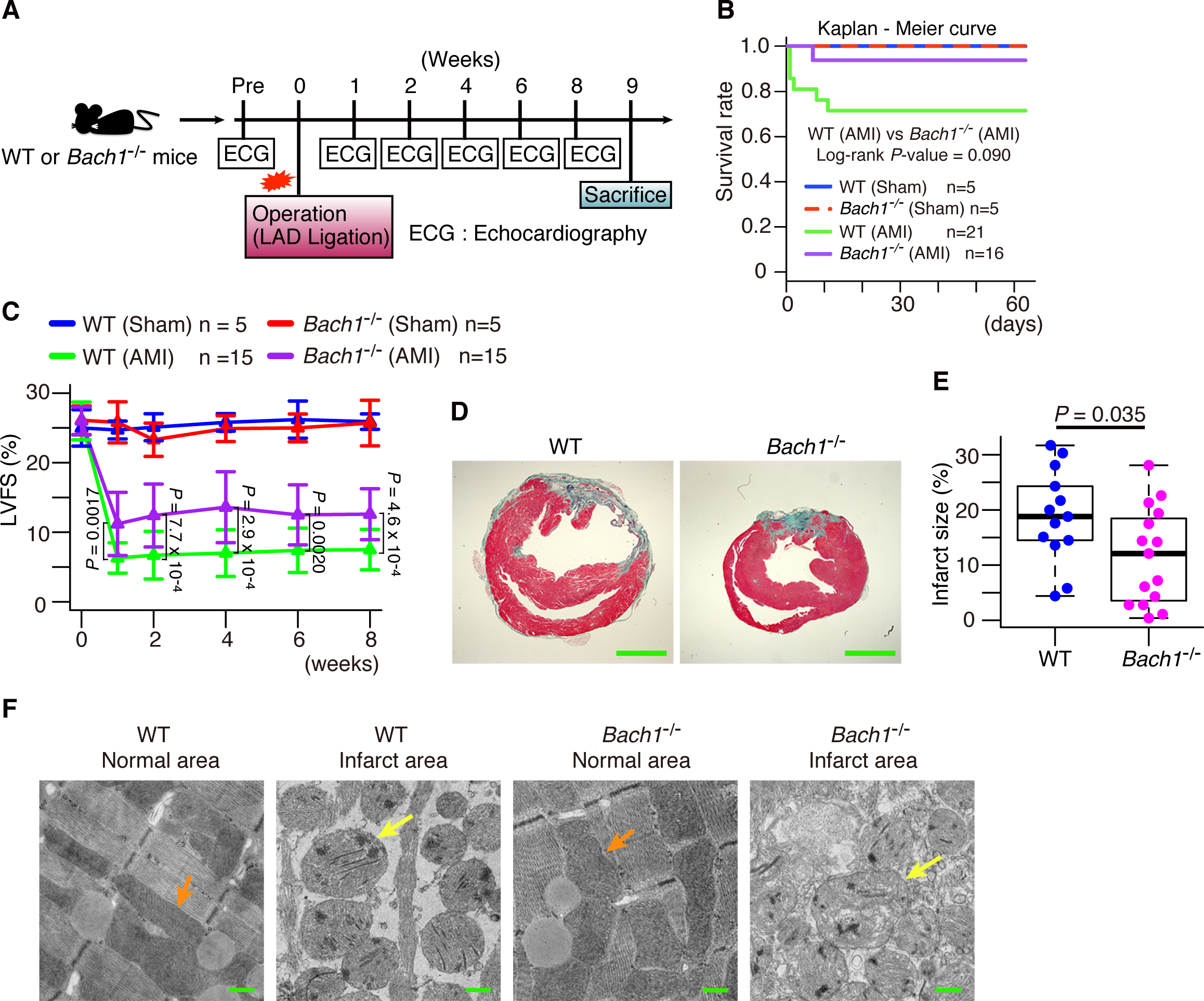
BACH1 aggravates AMI. A Experimental process. B Kaplan-Meier curve of each group. C Left ventricular fractional shortening (LVFS) on echocardiogram. D, E Mice was dissected after 9 weeks from operation. Representative photographs of heart sections stained with Elastica Masson staining (D). Infarct size to left ventricular section (E). Scale bars in (D) represent 2 mm. F Mice was dissected next day from operation. Transmission electron microscope image of normal and infarct area of hearts of mice next day from operation. Orange arrow: normal mitochondria. Yellow arrow: shrunken mitochondria. Scale bars represent 500 nm. Data information: Error bars of (C) represent standard deviation. The box and whisker plots of (E) show the 25th and 75th percentile quartiles and median values (center black line) and maximum and minimum values of the data. *P*-value of (B) by Log-rank test between WT (AMI) and *Bach1*^-/-^ (AMI). *P*-value of (C) by Tukey-Kramer method after two-way ANOVA. *P*-value of (E) by *t*-test.

In order to investigate whether or not ferroptosis is involved in the pathology, we observed the myocardial infarct regions using a transmission electron microscope. Shrunken mitochondria were observed in both WT and *Bach1*^-/-^ mice (Fig 6F). We then investigated whether or not the pathological changes could be improved by administering DFX, which is a clinically used iron chelator. First, we confirmed that it inhibited ferroptosis in MEFs (Figs 7A and EV5D, E). Although there was no improvement in the survival rates in WT or *Bach1*^-/-^ mice (Fig 7B), an improvement in the cardiac function on echocardiography was observed in the DFX group, which was more prominent in the WT mice than *Bach1*^-/-^ mice (Figs 7C, D and EV5F-K). The DFX group of WT mice showed a reduction in the infarct area; however, no such effect was noted in *Bach1*^-/-^ mice (Fig 7E and F). These results suggest that BACH1 exacerbates the pathology of AMI by promoting ferroptosis.

**Figure 7.**
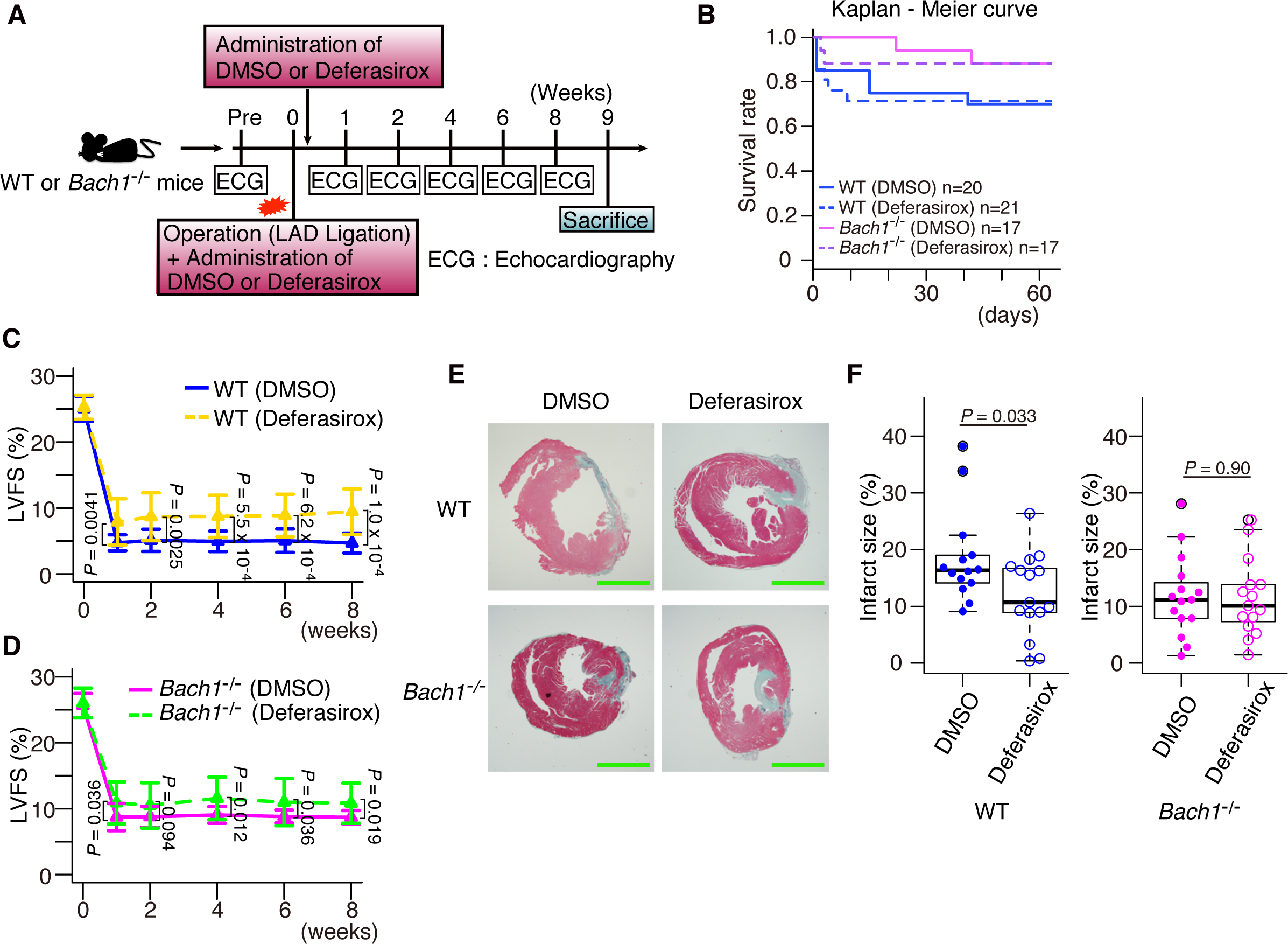
An iron chelator DFX alleviates AMI. A Experimental process. B Kaplan-Meier curve of each group. C, D Left ventricular fractional shortening (LVFS) of WT mice (C) and *Bach1*^-/-^ mice (D) on echocardiogram. E, F Mice was dissected after 9 weeks from operation. Representative photographs of heart sections stained with Elastica Masson staining (E). Infarct size to left ventricular section (F). Scale bars in (E) represent 2 mm. Data information: Error bars of (C, D) represent standard deviation. The box and whisker plots of (F) show the 25th and 75th percentile quartiles and median values (center black line) and maximum and minimum values of the data. *P*-value of (C, D, F) by *t*-test. Expanded View Figure 1. Transcription of *Bach1* and *Hmox1* increases in response to erastin. WT MEFs (7th, 9th, and 11th passage: P7, P9, and P11) were exposed to erastin for 10 hrs. qRT-PCR analysis for *Bach1* and *Hmox1* mRNA relative to *Actb* mRNA. Data information: Error bars represent standard deviation. *P*-value by Tukey’s test after one-way ANOVA.

## Discussion

While genes involved in ferroptosis are being discovered (Stockwell et al., 2017), how their expression is regulated during ferroptosis remains unclear. In this study, we found that many of the inhibitory genes of ferroptosis were coordinately upregulated upon induction of ferroptosis with erastin (Fig 1A). Such a coordinated response may be a mechanism for restricting ferroptosis. We further showed that BACH1 directly counteracted this coordinated response of genes, including *Hmox1, Slc7a11, Gclm, Gclc, Fth1, Ftl1*, and *Slc40a1* (Figs 3B, C and 5A, B), which are involved in the metabolism of GSH or labile iron. The protein amount of BACH1 was reduced upon the induction of ferroptosis (Fig 1B). Bach1 is known to repress the expression of *Slc40a1* in macrophages (Marro et al., 2010). Therefore, the reduction of BACH1 protein level may trigger the coordinated induction of these genes as a subprogram of the initial phase of ferroptosis program. Cells can then integrate distinct signals leading to BACH1 degradation, and thus judge whether or not they should undergo ferroptosis. Thus, BACH1 sets the threshold for whether or not ferroptosis occurs in response to lipid peroxide synthesized.

NRF2 is known to activate some of the genes that are repressed by BACH1, including *Hmox1*, *Slc7a11*, *Gclm* and *Gclc* (Alam et al., 1999, Bea et al., 2003, Ishii et al., 2000, Sasaki et al., 2002, Sekhar et al., 2003, Wild et al., 1999). Even though NRF2 increases the intracellular glutathione amount, it only weakly protects cells from ferroptosis (Cao et al., 2019). Other reports have shown that NRF2 can inhibit ferroptosis (Fan et al., 2017, Roh et al., 2017, Sun et al., 2016). Therefore, ferroptosis execution may depend on the initial amounts and kinetics of the induction or reduction of these transcription factors. This mechanism may extend our understanding of the regulation of ferroptosis, wherein ferroptosis is a cell death programmed at the level of the gene regulatory network.

We showed that GSH was higher in *Bach1*^-/-^ MEFs than WT cells (Fig 3A). Our results strongly suggest that BACH1 decreases intracellular GSH by repressing the expression of *Gclm*, *Gclc*, and *Slc7a11* (Fig 3B and C). Indeed, the protein amount of GCLM was higher in *Bach1*^-/-^ MEFs than in WT cells (Fig 3D). However, the protein amounts of GCLC and SLC7A11 were similar between WT and *Bach1*^-/-^ MEFs (Fig 3D). Cells may have additional mechanisms to tune strictly the protein amounts of GCLC and SLC7A11, managing the intracellular GSH amount and maintaining homeostasis. We found that SLC7A11 was further regulated by proteosomal degradation (Fig 3E). This observation suggests that the decision to undergo ferroptosis may be made based upon whether or not cells can induce efficiently inhibitory proteins like SLC7A11. Cells with higher amounts of SLC7A11 may likely be protected from ferroptosis. *Gclc* and *Slc7a11* may be critical factors for cells, with the transcriptional regulation by BACH1 and additional layers of regulation, although these points will need to be explored in further studies.

Reports on the function of HO-1 are conflicting, with studies conversely describing it to promote or inhibit ferroptosis (Adedoyin et al., 2018, Fang et al., 2019, Kwon et al., 2015, Sun et al., 2016). These discrepant findings may be due to the fact that HO-1 degrades prooxidant heme to produce not only the radical scavengers biliverdin and bilirubin but also free iron that mediates ferroptosis through Fenton reaction (Igarashi & Watanabe-Matsui, 2014, Stockwell et al., 2017). Therefore, in order to allow HO-1 to function effectively as an anti-oxidative stress enzyme, it is essential to suppress the reactivity of labile iron derived from heme. We showed that BACH1 represses the expression of the genes of ferritin and ferroportin (Fig 5A and B), which reduce the intracellular availability of labile iron. By increasing the expression of not only HO-1 but also ferritin and ferroportin during the induction of ferroptosis (Fig 1A), the prooxidant activities of heme and heme-derived free iron can be suppressed efficiently, thus protecting cells from ferroptosis. Conversely, BACH1 represses the expression of ferritin and ferroportin in addition to HO-1, thus effectively promoting ferroptosis (Fig 5C). Based on the present and previous findings, we proposed a model in which BACH1 accelerates ferroptosis by suppressing two major intracellular counteracting mechanisms against ferroptosis: the GSH synthesis pathway and the system for the sequestration of labile iron (Fig 5C).

In addition, we showed that ferroptosis was involved in the pathology of not only ischemia-reperfusion injury (IRI) (Baba et al., 2018, Fang et al., 2019, Gao et al., 2015) but also AMI. The severity of AMI was improved by the iron chelator, DFX particularly in WT mice (Fig 7C-F). The peripheral areas of AMI are naturally reperfused by angiogenesis, where ferroptosis is likely induced. Unexpectedly, DFX did not improve the survival rate. This may be explained by observations that adhesion between the cardiac infarct area and chest wall was smaller and cardiac rupture occurred more frequently in the DFX group than in the control group. These effects may offset the reduction in the infarct areas. Ferroptosis and subsequent inflammation may prevent cardiac rupture by pleural adhesion, but this issue needs to be investigated further. Nonetheless, our results here suggest that the therapeutic effect of DFX is expected in AMI and IRI. Necroptosis is also reportedly involved in cardiac ischemic disease (Oerlemans et al., 2012, Smith et al., 2007). Therefore, the double inhibition of ferroptosis and necroptosis may lead to the more effective palliation of AMI. In addition, this study suggests that *Bach1*^-/-^ mice are more resistant to AMI than WT mice because of their lower rate of ferroptosis than in WT mice (Figs 6 and 7). BACH1 may be a potential therapeutic target of AMI in the future.

Ferroptosis is thought to play a major role in cancer suppression (Jiang et al., 2015, Yang et al., 2014). Our results suggest that cancer cells may acquire resistance against ferroptosis by decreasing BACH1 protein, thus eluding elimination by ferroptosis. We previously reported that BACH1 promotes the proliferation of MEFs transformed with H-Ras^v12^ and their tumor formation in a mouse transplantation model (Nakanome et al., 2013). Recently, BACH1 was found to promote the proliferation and/or metastasis of breast cancer and ovarian cancer cells (Han et al., 2019, Lee et al., 2014, Lee et al., 2019, Mansoori et al., 2019). BACH1 is therefore considered to have dual functions in cancers: promoting cell proliferation and cell death through ferroptosis. Cancer cells may adapt to their surrounding environment by changing the expression of BACH1; cancer cells may highly express BACH1 during stages of proliferation and metastasis but may reduce their levels of BACH1 under stress conditions, such as toxicity due to anti-cancer drugs. Such flexibility in the amount of BACH1 protein expressed may enhance the malignancy of cancer cells. Therapy that targets this flexibility, such as the down-regulation of BACH1 in response to erastin, may expand the field of potential cancer treatments.

## Materials and Methods

### Mice

The generation of *Bach1*^−/−^ mice on the C57BL/6J background was described previously (Sun et al., 2002). Mice 13 weeks of age were analyzed for models of AMI. Animals were euthanized by cervical dislocation under anesthetic inhalation overdose with isoflurane before anatomy. These mice were bred at the animal facility of Tohoku University. Mice were housed under specific pathogen-free conditions. All experiments performed in this study were approved by the Institutional Animal Care and Use Committee of the Tohoku University Environmental & Safety Committee.

### Mice models of AMI

Induction of AMI was performed as described previously (Abarbanell et al., 2010, Shindo et al., 2016). The mice were subjected to ligation of the proximal left anterior descending coronary artery (LAD) to induce AMI. They were randomly assigned to sham or AMI group (Fig 6A), DMSO or DFX group (Fig 7A). In order to follow up the time course of LV function after AMI, we performed transthoracic two-dimensional echocardiography. For histological analysis and analysis with transmission electron microscope, the heart was divided along the short axis at the center of the infarct.

### Histopathological Analysis

Excised hearts were fixed with 4% paraformaldehyde for histological and immunohistochemical examination. After 24-48 hours of fixation and dehydration through increasing concentrations of ethanol, the tissue specimens were embedded in paraffin and sliced at 3 µm in thickness. The sections were used for Elastica-Masson staining. The extent of infarct area was calculated as a rate of fibrotic area using the following formula: fibrotic area / (LV free wall + interventricular septum) x 100 (%) with use of Photoshop software (Adobe).

### Transmission electron microscopy

Cells and hearts were treated in 2.5% glutaraldehyde in 0.1 M Cacodylate buffer [pH 7.4] for at least 24 hrs, and washed with 0.1 M Cacodylate buffer 4 times and then treated with 1% OsO4 in 0.1 M Cacodylate buffer for 90 min. After dehydration through an ethanol series (50-100% ethanol), cells were embedded in Epon resin. Thin sections were cut with a microtome (Leica EM UC-7), stained with 2% uranyl acetate and 0.4% lead citrate, and examined and photographed under a transmission electron microscope (Hitachi High-Technologies H-7600).

### Isolation and culture of MEFs

MEFs were derived from 13.5-day-old embryos of WT or *Bach1*^-/-^ mice. Following removal of the head and organs, embryos were rinsed with PBS (Nissui, Tokyo, Japan), minced and digested with trypsin (0.05% (v/v) solution containing 0.53 mM EDTA) (Gibco, Carlsbad, CA, USA) and 1.8 mg/ml DNase I (Roche, Basel, Switzerland) in PBS and incubated for 60 min at 37°C. Trypsin was inactivated by addition of DMEM with high glucose (Gibco) containing 10% (v/v) fetal bovine serum (FBS) (Sigma-Aldrich, St. Louis, MO, USA), 1x MEM nonessential amino acids (Gibco), and 0.1 mM 2-mercaptoethanol (Sigma-Aldrich)). MEFs from a single embryo were plated into a 100-mm diameter culture dish and incubated at 37°C in 3% oxygen (1st passage: P1). MEFs from embryos of homosexual littermates were mixed at 2nd passage (P2) and stocked.

MEFs were maintained at 37°C in culture medium (DMEM with (Gibco) containing 10% FBS (Sigma-Aldrich), 1x MEM nonessential amino acids (Gibco), penicillin/streptomycin (100 U/ml and 100 µg/ml each) (Gibco) and 0.1 mM 2-mercaptoethanol (Sigma-Aldrich)) in 3% oxygen for experiments. The number of passage were recorded for each rot of MEFs. From 5th to 11th passage MEFs were used for all experiments.

### Reagents

Erastin, DMSO, and DFO were purchased from Sigma-Aldrich. MG132 was purchased from Calbiochem (San Diego, CA, USA). DFX was transferred as raw material from Novartis Pfarma (Basel, Switzerland). S-adenosylmethionine (SAM) -^13^C_5_, ^15^N and Glutathione (GSH) -^13^C_2_, ^15^N were purchased from Taiyo Nissan Corp. (Tokyo, Japan) and used as internal standard (IS) for mass spectrometry. Methanol, acetonitrile and ammonium hydroxide for mass spectrometry were purchased from Kanto Chemical (Tokyo, Japan). Ammonium bicarbonate (1 mol/L) for mass spectrometry was purchased from Cell Science & Technology Inst., Inc. (Miyagi, Japan). Formic acid for mass spectrometry was purchased from Wako Pure Chemical Industries (Osaka, Japan).

### Sample preparation for UHPLC/MS/MS

MEFs (3-8 x 106 cells for each lot) were suspended in 100 µL of methanol containing the internal standards (0.2 µg/mL SAM-13C515N for positive ion mode (Pos) and 1 µg/mL GSH-13C215N for negative ion mode (Neg)), and were homogenized by mixing for 30 sec followed by sonication for 10 min. After centrifugation at 16,400 x g for 20 min at 4°C followed by deproteinization, 3 µL of each extract was analyzed by ultra high-performance liquid chromatography triple quadrupole mass spectrometry (UHPLC/MS/MS).

### UHPLC/MS/MS analysis

The UHPLC/MS/MS analysis was performed on an Acquity™ Ultra Performance LC I-class system (Waters Corp. Milford, UK) interfaced to a Waters Xevo TQ-S MS/MS system equipped with electrospray ionization (ESI). The MS/MS was performed using the multiple reaction monitoring (MRM) mode with a scan time of 1 ms for each compound. The transitions of the precursor ion to the product ion, cone voltage and collision energy are listed in Appendix Table S1. The other settings are as follows: 3.5 kV (Pos) or 2.5 kV (Neg) capillary voltage, 30 V cone voltage, 50 V source offset, source temperature at 150°C, 150 L/hr cone gas (N2) flow rate, desolvation temperature at 450°C, 1000 L/hr desolvation gas flow, 0.15 min/mL collision gas flow, 7.00 bar nebulization gas (N_2_) flow. LC separation, was performed as described before (Saigusa et al., 2016), using a normal-phase column (ZIC-pHILIC; 100 mm × 2.1 mm i.d., 5 µm particle size; Sequant, Darmstadt, Germany) with a gradient elution using solvent A (10 mmol/L NH_4_HCO_3_, adjusted to pH 9.2 using ammonia solution) and B (acetonitrile) at 300 µL/min: 99 to 70% B from 0.5 to 4.0 min, 70 to 1% B from 4.0 to 6.5 min, 1% B for 2.5 min, and 99% B for 9 min until the end of the run. The oven temperature was 20°C. The data were collected using the MassLynx v4.1 software (Waters Corp.) and the ratio of the peak area of analyte to the IS was analyzed by Traverse MS (Reifycs Inc., Tokyo, Japan).

### RNA interferrence

All siRNAs (siControl: Stealth RNAi^TM^ siRNA Negative Control, Med GC, siGclm #1: MSS204722, siGclm#3: MSS204724, siSlc7a11 #1: MSS218649, siSlc7a11 #2: MSS218650, siGclc #2: MSS204720, siGclc #3: MSS204721, siHmox1 #1: MSS247281, siHmox1 #3: MSS274857) were obtained from Invitrogen (Carlsbad, CA, USA). Sequences of the siRNAs are described in Appendix Table S2. 2 x 10^6^ cells of MEFs were transfected with 1.2 nM of siRNAs using Amaxa Nucleofector II (Lonza, Basel, Switzerland) and MF 1 Nucleofector kit (Lonza) according to the manufactures protocols. After transfection, MEFs were passaged to dishes or culture plate with culture medium.

### Western Blotting

Cells were trypsinized, pelleted, and washed twice in PBS. Cells were lysed beyond 5 min in SDS sample buffer (62.5 mM Tris-HCl (pH = 6.8), 1% (v/v) 2-Mercaptoethanol、1% (w/v) Sodium dodecyl sulfate; SDS, 10% (w/v) Glycerol, 0.02% (w/v) Bromophenol blue; BPB). Lysates were resolved on 7.5–10% SDS–PAGE gels and transferred to PVDF membranes (Millipore, Billerica, MA, USA). The antibody for detection of β-actin (sc-1616) was purchased from Santa Cruz Biotech (Santa Cruz, CA, USA). The antibody for detection of HO-1 (ADI-SPA-896) was purchased from Enzo life science (New York, NY, USA). The antibodies for GAPDH (ab8245), Gclc (ab53179), and Gclm (ab124827) were purchased from Abcam (Cambridge, UK). The antibody for Slc7a11 (119-11215) was purchased by RayBiotech (Norcross, GA, USA). The antibody for BACH1 was described previously (Sun et al., 2002).

### Quantitative PCR with reverse transcription

Total RNA was purified with RNeasy plus micro kit or RNeasy plus mini kit (Qiagen, Hilden, Dermany). Complementary DNA was synthesized by a SuperScript III First-Strand Synthesis System (Invitrogen). Quantitative PCR was performed using LightCycler Fast Start DNA Master SYBR Green I, and LightCycler nano (Roche) or LightCycler 96 (Roche). mRNA transcript abundance was normalized to that of Actb. Sequences of the qPCR primers are described in Appendix Table S3.

### Administration of erastin and Cell death assessment by flow cytometry

Before administration of erastin, the medium was exchanged to the experimental medium (culture medium without 2-mercaptoethanol and penicillin/streptomycin). Erastin was dissolved in DMSO and adiministered to experimental medium with DMSO. The concentration of DMSO was adjusted among each samples. Cell death was assessed 24 hours after administration of erastin. PI and Annexin V staining were used for assessment of cell death. APC-Annexin V was purchased from Becton, Dickinson and Company (BD) (Franklin Lakes, NJ, USA). MEFs were stained by APC-Annexin V according to the manufactures protocols. PI was added to aliquot (1 µg/mL) before flow cytometry. The MEFs were sorted with a FACS Aria II (BD) and analyzed by FlowJo software (Tree Star, Ashland, OR, USA). MEFs of positive of whether at least Annexin V or PI was assessed as dead cells. The gating strategy for assessing dead cells (Figs 2B, E, 4B-E, and EV5E) was shown in Fig EV2.

### ChIP-Seq

We used ChIP-seq data of BACH1 and MAFK in M1 cell line from GEO (Gene Expression Ominibus) data set GSE79139 that deposited for our previous report (Ebina-Shibuya et al., 2017, Ebina-Shibuya et al., 2016).

### RNA-Seq

Total RNA was purified using an RNeasy plus mini kit (Qiagen). To remove ribosomal RNA (rRNA), 4 µg of the total RNA was treated with a GeneRead rRNA Depletion kit (Qiagen) and then with an RNeasy MiniElute kit (Qiagen) for cleanup. For fragmentation, 100 ng of the rRNA-depleted RNA was incubated at 95°C for 10 min and was purified by a Magnetic Beads Cleanup Module (Thermo Fisher Scientific, Carlsbad, CA, USA). The libraries were constructed with an RNA-seq library kit ver. 2 (Thermo Fisher Scientific) on ABI library builder (Thermo Fisher Scientific), and was barcoded with Ion Xpress RNA-seq BC primer (Thermo Fisher Scientific). The library fragments with a size range of 100-200 bp were selected with Agencourt AMPure XP beads (Beckman Coulter, Brea, CA, USA). Templates were prepared on the Ion Chef system using an Ion PI Hi-Q Chef kit (Thermo Fisher Scientific) and sequencing was performed on an Ion Proton system using with Ion PI Hi-Q sequencing kit (Thermo Fisher Scientific) the PI v3 chip (Thermo Fisher Scientific). The sequence data were obtained as fastq files. The sequence data was aligned to reference hg19 using the RNASeqAnalysis plugin from Ion torrent suite software (Thermo Fisher Scientific). Mapped reads were counted for each gene using HTSeq v 0.9.1 htseq-count. The differential expression analysis was performed on edge R v 3.16.5 after removal of low count lead genes using three biological replicates for each condition (less than 5 leads per gene in the sample and counts per million mapped reads (CPM) of 1 or less).

### Statistics

For all experiments, differences of data sets were considered statistically significant when *P*-values were lower than 0.05. Statistical comparisons were performed using the *t*-test in comparison between the two groups, and one, two, or three way ANOVA followed by Tukey’s test or Tukey-Kramor method in comparison among multiple groups. For the *t*-test, student’s *t*-test was used when the standard deviation (SD) of the groups was not significantly different by *f*-test. Welch’s *t*-test was used when the SD of the groups was significantly different by *f*-test.

## Data Availability

The RNA-seq data has been deposited at the GEO database under accession codes GSE131444.

## Acknowledgements

We thank members of the Departments of Biochemistry, Tohoku University Graduate School of Medicine for discussions and support; the Biomedical Research Core of Tohoku University Graduate School of Medicine for technical support. We thank Novartis for raw material transfer of DFX for this study. This study was supported in part by Grants-in-Aid from the Japan Society for the Promotion of Science (19K07680 to M.M. and 15H02506, 24390066, 21249014 and 18H04021 to K.I.) and Agency for Medical Research and Development (JP15gm0510001 to K.I.).

## Author Contributions

Writing of original draft; H.N. Conceptualization and methodology; H.N., M.M and K.I. Major investigation; H.N. Bioinformatics analysis; H.N. and M.M. Advise and support for mice AMI model; T.S. and H.S. Supportive investigation; H.K., K.S., M.S., and Y.I. Investigation of UHPLC/MS/MS; D.S. Review and editing; H.N and K.I. Supervision; K.I.

---

**Expanded View Figure 1.**
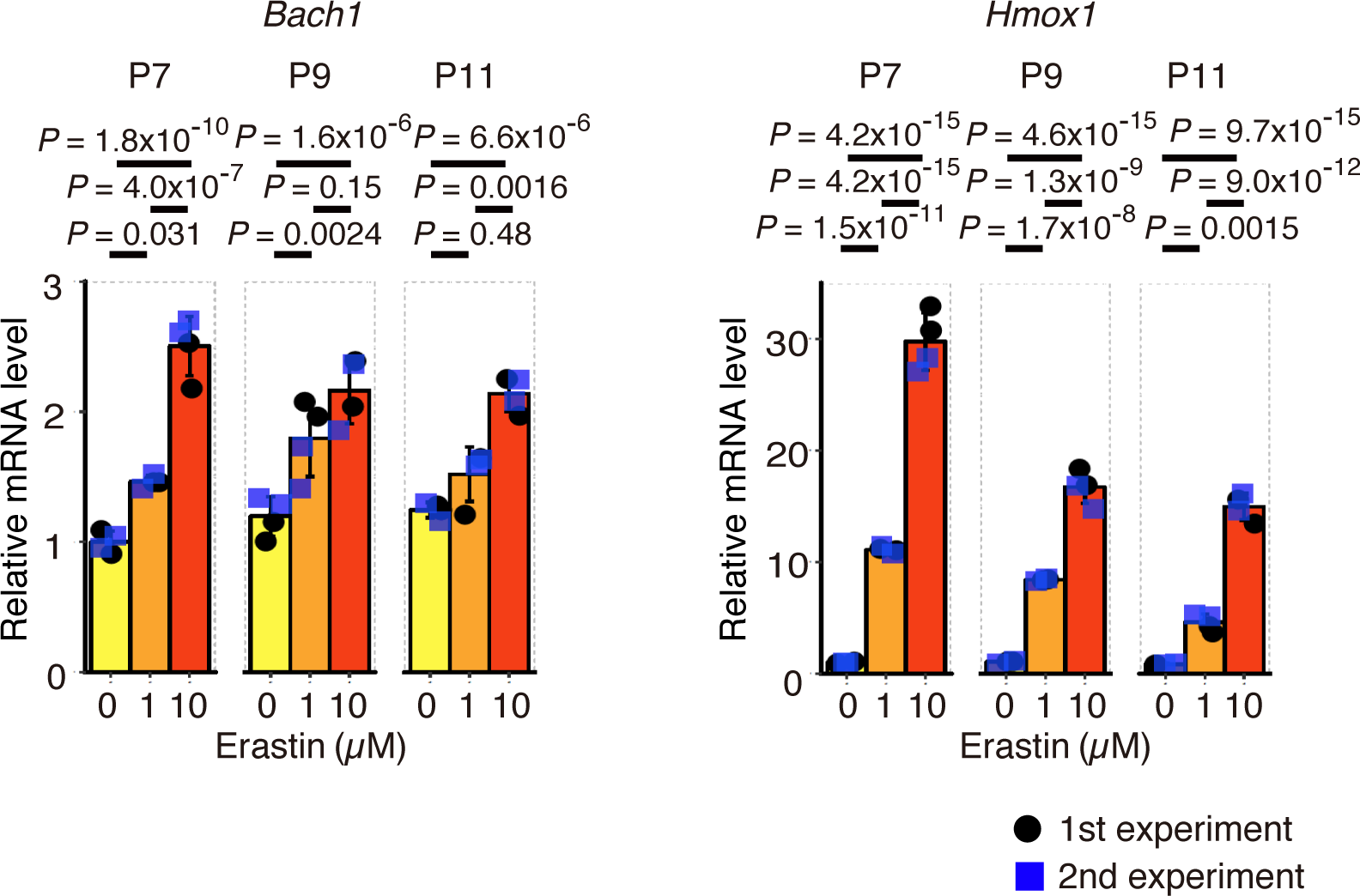
Transcription of *Bach1* and *Hmox1* increases in response to erastin. WT MEFs (7th, 9th, and 11th passage: P7, P9, and P11) were exposed to erastin for 10 hrs. qRT-PCR analysis for *Bach1* and *Hmox1* mRNA relative to *Actb* mRNA. Data information: Error bars represent standard deviation. *P*-value by Tukey’s test after one-way ANOVA.

**Expanded View Figure 2.**
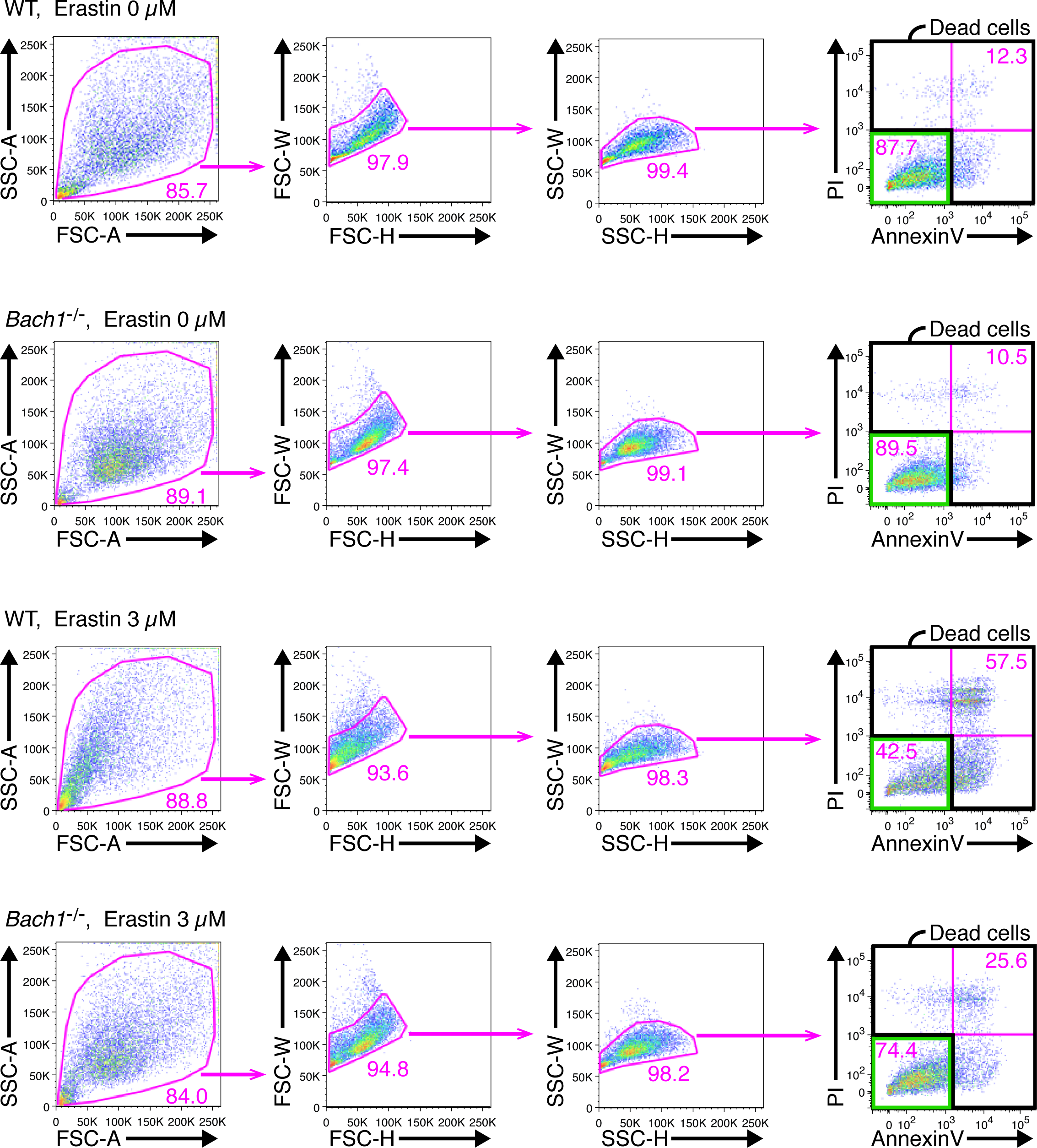
Additional data demonstrating flow cytometry gating of dead cells in WT and *Bach1*^-/-^ MEFs exposed to erastin. WT and *Bach1^-/-^*MEFs (11th passage: P11) exposed to erastin for 24 hrs (Figure 2B). Representative flow cytometry images showing the strategy that was implemented for the sorting of dead cells. Propidium iodide (PI) positive or Annexin V positive cells were judged as dead cells. The similar strategy was implemented in Figs 2E, 4B-E, and EV5E.

**Expanded View Figure 3.**
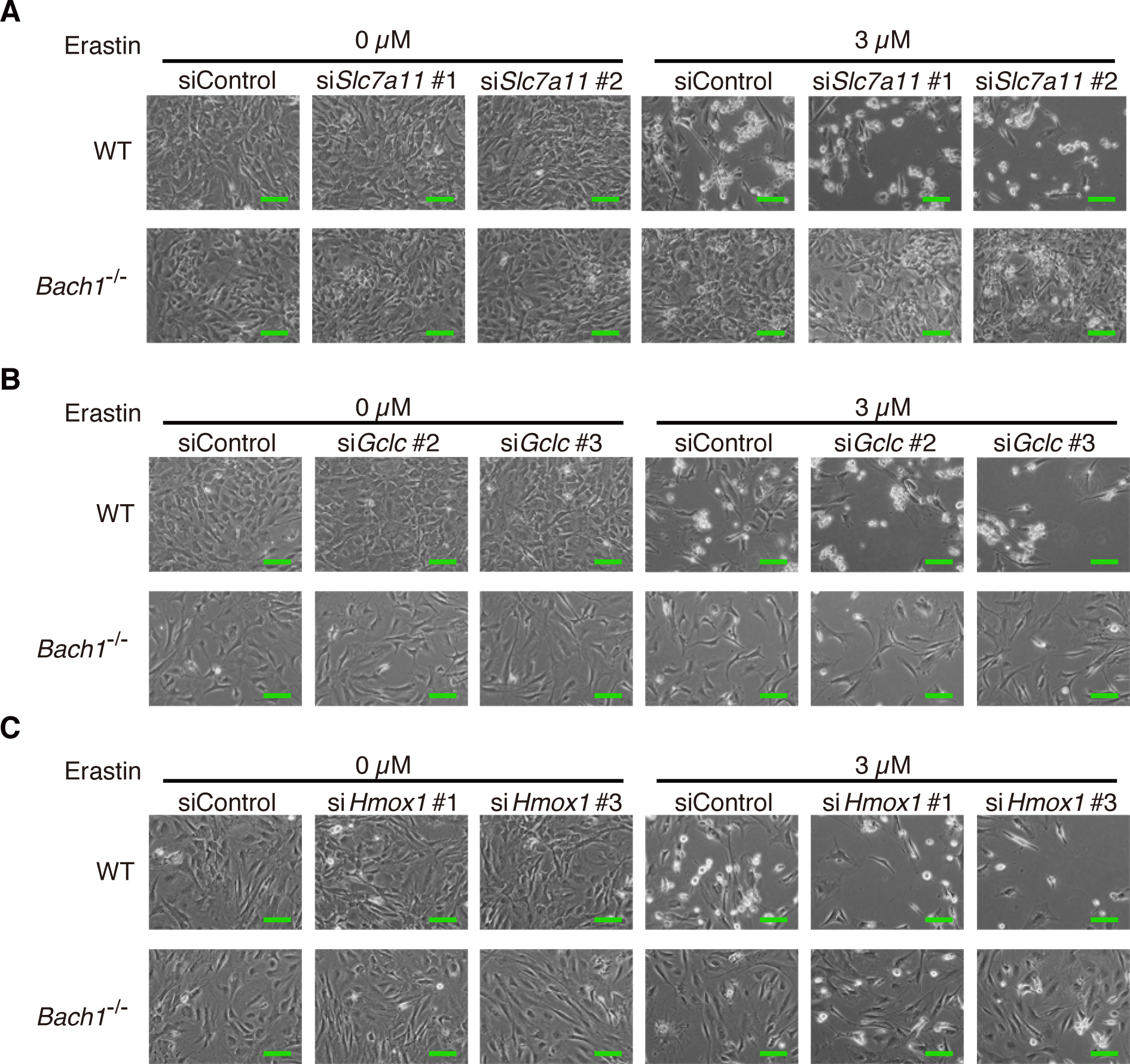
*Slc7a11*, *Gclc*, and *Hmox1* repress ferroptosis, associated with **Fig 4**. A-C siRNA was transfected to WT and *Bach1*^-/-^ MEFs (5th or 6th passage). After 24 hrs, MEFs were exposed to erastin for 24 hrs. Optical microscope image. Scale bars represent 100 µm.

**Expanded View Figure 4.**
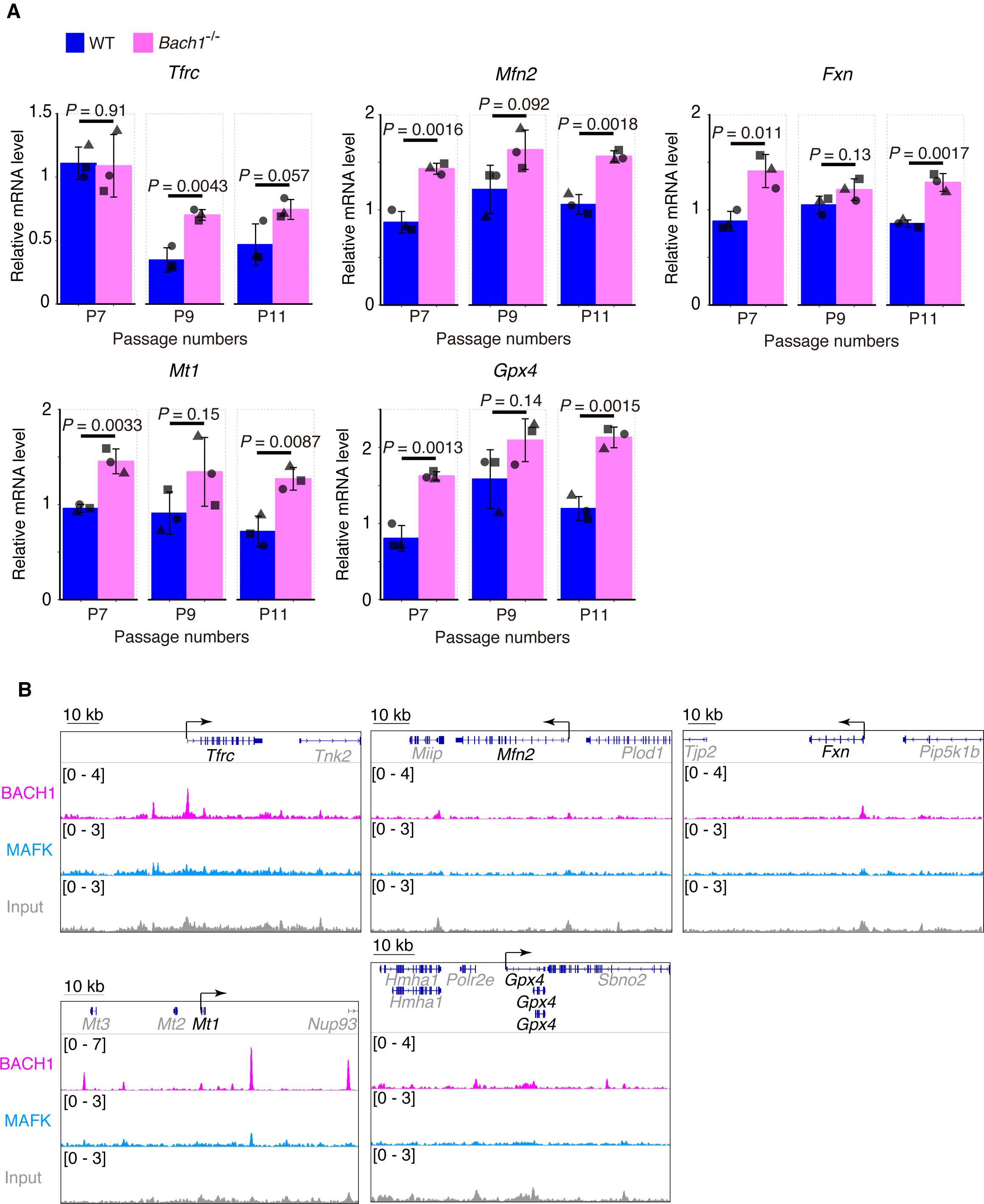
*Tfrc*, *Mfn2*, *Fxn*, *Mt1*, and *Gpx4* were not strongly regulated by BACH1. A qRT-PCR analysis for *Tfrc*, *Mfn2*, *Fxn*, *Mt1*, and *Gpx4* mRNA relative to *Actb* mRNA in WT and *Bach1^-/-^* MEFs (7th, 9th, and 11th passage: P7, P9, and P11). B ChIP-seq analysis of the binding of BACH1, MAFK for gene regions of *Tfrc*, *Mfn2*, *mFxn*, *Mt1*, and *Gpx4* in M1 cells. Data information: Error bars of (A) represent standard deviation. *P*-value of (A) by *t*-test.

**Expanded View Figure 5.**
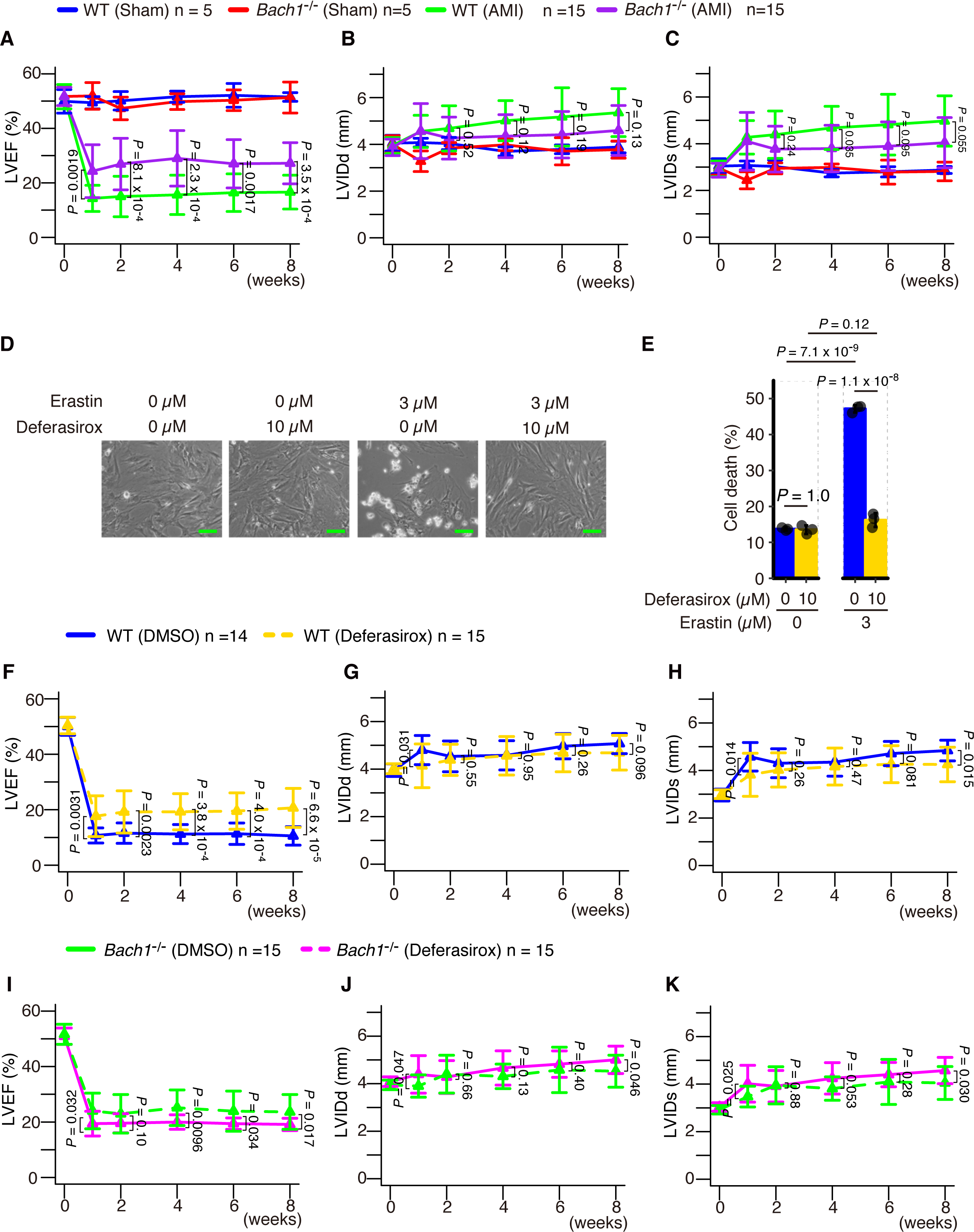
BACH1 aggravates AMI, that was alleviated by an iron chelator DFX. A-C Left ventricular ejection fraction (LVEF) (A), left ventricular internal dimension in diastole (LVIDd) (B), and left ventricular internal dimension in systole (LVIDs) (C) on echocardiogram. D, E Optical microscope image (D) and Quantification of cell death by flow cytometer (E) of WT and *Bach1^-/-^* MEFs (10th passage: P10) exposed to erastin for 24 hrs. Scale bars in (D) represent 100 µm. (F-K) Left ventricular ejection fraction (LVEF) (F, I), left ventricular internal dimension in diastole (LVIDd) (G, J), and left ventricular internal dimension in systole (LVIDs) (H, K) on echocardiogram. Data information: Error bars of (A-C, E-K) represent standard deviation. *P*-value of (A-C) by Tukey-Kramer method after two-way ANOVA. *P*-value of (E) by Tukey’s test after two-way ANOVA. *P*-value of (F-K) by *t*-test.

